# Marker-independent imaging reveals a correlation of fibrotic and epigenetic alterations in endometriosis

**DOI:** 10.1101/2025.07.10.664232

**Authors:** Tara Beyer, Lucas Becker, Sema Demir, Simone Liebscher, Daniel Carvajal-Berrio, Hans Bösmüller, Katharina Rall, Bernhard Krämer, Sara Y Brucker, Katja Schenke-Layland, Martin Weiss, Julia Marzi

## Abstract

**Introduction:** Endometriosis describes the presence of endometrial glands outside of the uterus and can cause various symptoms such as chronic pelvic pain, hypermenorrhea and infertility. These complications pose an extreme burden on the patients, especially as up to date, the average time until diagnosis can consume several years and requires invasive laparoscopy.

**Objectives:** The aim of this study is to molecularly characterize endometrium and endometriosis using marker-independent Raman microspectroscopy to identify potential biomarkers and validate its diagnostic potential.

**Methods:** After histopathological characterization of tissue sections of human endometrium and endometriosis, Raman microspectroscopy was performed on the gland region. Multivariate analysis of the hyperspectral maps was used to localize major subcellular structures and further decipher their molecular composition. Samples from different anatomical regions and throughout all menstrual cycle phases were analyzed.

**Results:** Raman imaging enabled label-free visualization of tissue morphology and submolecular tissue characterization. Distinct differences between endometrium and endometriosis were found for collagen type I and nuclear signatures. Spectral deconvolution allowed identification of a Raman biomarker indicative of fibrotic changes in endometriosis samples. Additionally, a significant increase in epigenetic 5mC foci and an increased signal intensity relevant for methylations was detected in nuclei of endometriosis. Furthermore, a neural network-based classification of Raman data resulted in high accuracies in discriminating endometrial and peritoneal tissue from endometriosis.

**Conclusion:** The non-destructive approach by hyperspectral Raman imaging enabled for molecular sensitive characterization of endometriotic lesions which could not only be an asset in complementing histopathological tissue evaluation but combined with data-driven classification models support in situ tissue diagnosis.

## Introduction

Endometriosis is a benign, hormonally dependent disease affecting 10-15% of women in childbearing age; although the number of unreported cases is estimated to be much higher (1, 2). The severity of the disease varies from patient to patient. While about 50-60% of the affected women suffer from symptoms like dysmenorrhea, dyspareunia, dysuria, dyschezia, pelvic or bowel pain and abdominal bloating others are not aware of their condition until they encounter infertility disorder (3, 4). The determining feature of endometriosis is the appearance of endometrial tissue, which includes glands and stroma at an ectopic site, outside the uterus (5). Due to the unclear pathogenesis and heterogenous symptoms, there remains a lack of sensitive biomarkers for investigating and diagnosing endometriosis. As a result, invasive procedures like laparoscopy and histopathological biopsy continue to be the gold standard for diagnosis. The average diagnostic delay in endometriosis is approximately seven years.

One major obstacle is the indeterminate pathogenesis of endometriosis. Hitherto, there are several theories but none of them fully captures the manifold clinical appearances and disease patterns (6). It is likely that there is either a combination of the theories or other factors involved, such as inflammatory processes, hormone levels or genetic predisposition, which might play an important, yet undiscovered role (6, 7). Sampson’s theory about retrograde menstruation and consecutive ectopic implantation is the most supported in literature until today (8, 9). In the genetic-epigenetic theory, endometriosis is not solely based on genetic changes, but is at least partially epigenetically determined (10). Examples of causative factors are oxidative stress in the uterus during menstruation and in the abdominal cavity after the onset of retrograde menstruation, inflammation or immunological influences (11). This theory is consistent with all observations in endometriosis and would also explain that lesions respond to estrogen, progestins, and pregnancy (12, 13). Glands in ectopic locations can remain inactive for long periods without alteration (14).

Tissue from patients suffering from endometriosis has also shown features of fibrosis (5, 15, 16), which plays a pivotal role in the progression and worsening of the symptoms due to stiff lesions distorting the uterine anatomy (15, 17). This has resulted in a proposed revision of the definition and potential therapeutic targets of endometriosis, suggesting to focus on the fibrotic pathology (18, 19).

Furthermore, the last decade witnessed advances in non-destructive imaging techniques providing spatially and temporally resolved insights into tissue alterations. Besides clinical imaging such as transvaginal ultrasound, magnetic resonance imaging or computed tomography scans, novel, marker-independent imaging techniques gain relevance in complementary histopathological analyses and tissue discrimination (20, 21). Raman microspectroscopy (RMS) enables marker-independent and molecular-sensitive identification and localization of subcellular structures by analyzing molecular fingerprints of Raman-active biomolecules such as lipids, collagens, or nucleic acids reflecting a specific tissue state or cellular phenotype (22, 23, 24).

In this study, fibrotic and epigenetic features of endometrium and endometrium-like tissues from two anatomical regions - the ligamentum sacrouterinum (LiS) and the septum rectovaginale (SRV) - were identified utilizing conventional histology and characterized by complementary non-destructive RMS. The objective of this study was to gain a deeper understanding of modifications in pathological features in endometriotic glands and to identify endometriosis-relevant biomarkers. RMS was combined with machine learning-based techniques to recognize disease pathomechanisms and to develop a multiparametric data-driven model capable of differentiating between normal and diseased tissue stages throughout all phases of the menstrual cycle (Figure 1a).

**Figure 1.**
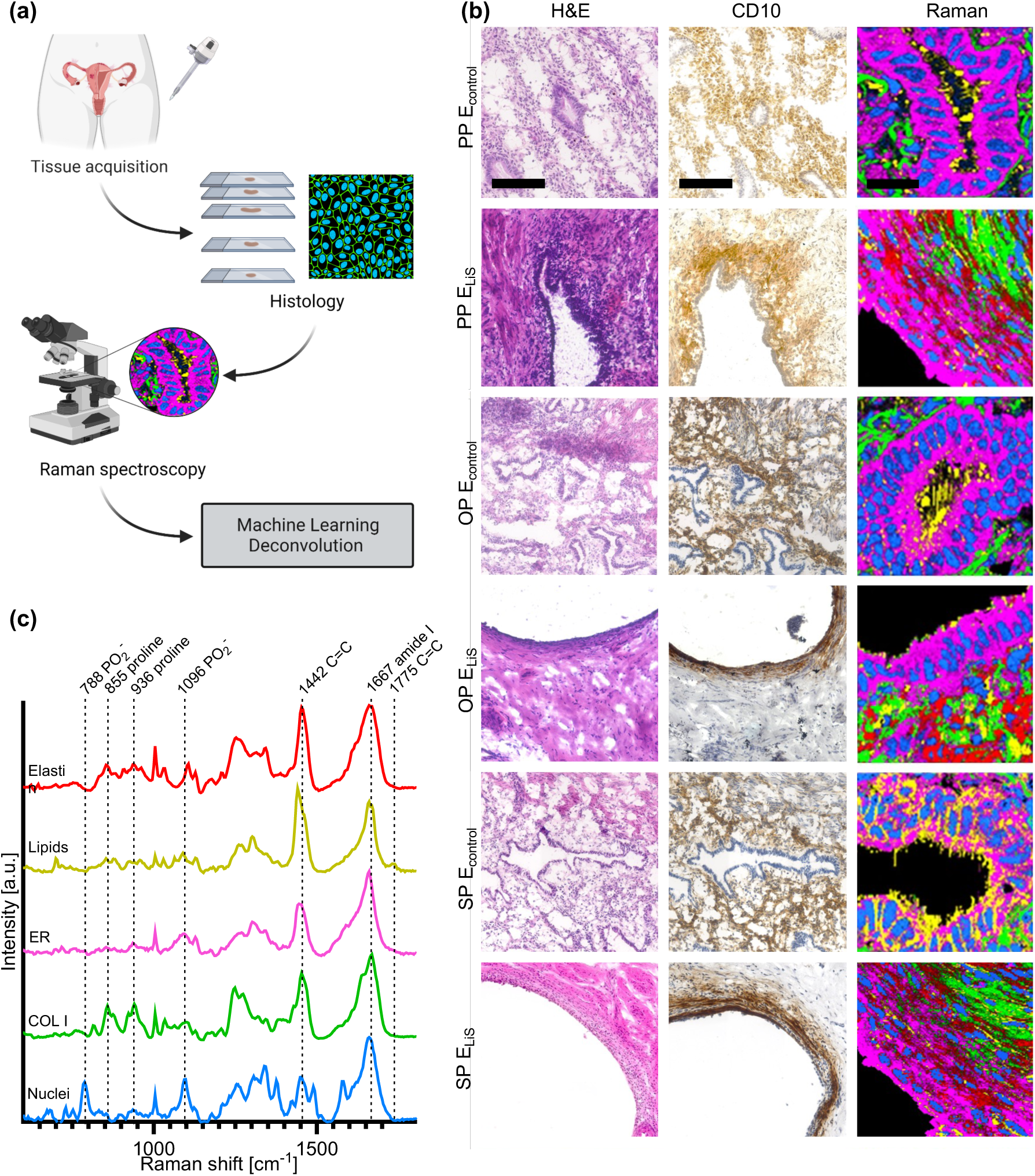
Conventional and Raman-based histopathology of endometrium and endometriotic glands in different menstrual cycle phases. (a) Workflow of histopathological and Raman spectroscopy-based diagnosis of endometrium and endometrial lesions. (b) Histology of endometrium (E_control_) and endometriosis from ligamentum sacrouterinum (E_LiS_) in proliferative phase (PP), ovulation phase (OP) and secretory phase (SP). H&E and CD10 of serial sections display endometrial glands and surrounding stromal cells. Scale bar equals 130 µm. Raman images of glands generated by true component analysis of Raman heatmaps. Scale bar equals 20 µm. (c) Corresponding Raman spectra identified by TCA: nuclei (blue), collagen type I (COL I, green), endoplasmic reticulum (ER, pink), lipids (yellow), elastin (red).

## Material and Methods

### Ethics statement

After informed consent was obtained, endometrium, endometriosis and peritoneal tissues were retrieved from surgical specimens. The Ethical Committee of the Medical Faculty of the Eberhard Karls University Tübingen approved the scientific use of the tissue (152/2018BO2).

### Collection of human tissue samples

Samples of endometrium, endometriosis from the LiS and the SRV, as well as peritoneum were acquired by the Women’s Hospital at the University Hospital Tübingen. Endometrium was procured through hysterectomy, while the endometriosis tissues were obtained via minimally invasive laparoscopic surgery. Only patients that were not taking contraceptives and had not yet reached menopause were included in the study. Specimens were stored in DPBS until cryo-preservation using Tissue Tek O.C.T. (Tissue Tec, Sakura Finetek Europe B.V., Alphen aan den Rijn, Netherlands). Serial 10 µm thick cross sections of endometrium, endometriosis and peritoneum tissue were prepared utilizing a cryotome (MICROM HM560, Thermo Scientific, Waltham, USA). Tissue sections were collected on objective slides (SUPERFROST PLUS, Menzel, Braunschweig, Germany) for histochemical and immunofluorescence staining, as well as Raman microspectroscopy. Overall, female patients with and without endometriosis (LiS) in all menstrual cycle phases (proliferative phase (PP), ovulation phase (OP) and secretory phase (SP)) were included (*n*=3). Additionally, SRV endometriosis and peritoneum were acquired (n=3). An overview of patient specific information is given in Supplementary Table S1.

### Histological Tissue Characterization

For characterization of cryosections from endometrium, endometriosis and peritoneum, routine immunohistochemical stains were performed. To determine general cellular and tissue morphology, cryosections were stained with hematoxylin-eosin (H&E) according to standard protocols. Consecutive CD10 staining served as confirmation stain to identify the localization of endometriosis. Ki67 staining was implemented to identify the proliferation of cells, while resorcin-fuchsin staining identified microfibrils and elastic fibers in tissues. Picrosirius-red (PSR) staining was implemented to visualize collagen maturity/thickness. Detailed protocols are available in the supplementary methods.

### 5mC Staining

Immunofluorescence (IF) 5mC staining was adapted from Daum et al (25) using a 5mC mouse monoclonal IgG antibody (Merck, MABE146, 1:2000, stock 2 mg/ml) at 4°C overnight and a goat anti-mouse IgG Alexa Fluor 488 (1:250, Thermo Fisher Scientific) secondary antibody.

### Brightfield and Immunofluorescence Imaging

Brightfield tile scans of H&E, CD10, resorcin-fuchsin and Ki67 stained tissue sections were performed on an Axio Observer using a 63x/1.4 NA C-plan apochromat objective (Zeiss Microscopy GmbH).

### 3D image acquisision and analysis

IF stains were imaged utilizing a Zeiss LSM 880 (Zeiss Microscopy GmbH). Images were acquired as z-stacks with 2 tracks through a 63x/1.4 NA C-plan apochromat objective. Z-stack images were analyzed with Imaris 9.7.2. Detailed information of image processing can be found in the supplementary methods.

### Raman Imaging

Raman scans were acquired on a customized setup as previously described (26). For each sample, glandular regions were measured according to ROIs defined by CD10 staining. Three spectral maps containing one spectrum per 0.5 µm were generated of an area of 80×80 µm, at an acquisition time of 0.05 s per spectrum and a laser power of 50 mW.

### Spectral Analyses

Image analyses of spectral maps were performed with the Project Five 5.2 software (WITec GmbH). Raman data were subjected to cosmic ray removal, polynomial baseline correction, cropped to 400 - 3000 cm^-1^ and area normalized to 1 prior to analysis by true component analysis (TCA). TCA identified spectral components that were most prevalent in the data set and enabled a visualization based on the generation of intensity distribution images for each identified component. Gray value intensities were determined in the intensity distribution images for quantitative assessment of each component in endometrium and endometriosis. Sum-filter images at the spectral position 1370 cm^-1^ a spectral width of 30 cm^-1^ were extracted from nuclei Raman spectra and combined with filter images recorded at 788 cm^-1^ depicting the whole nuclei. For image combination, the color top-value of the 5mC sum filter image was set to 0.1 with a bottom-value of 0, while the top-value of the nuclei filter image was set to 1 with a bottom-value of 0. Next, combined images were transferred to ImageJ 1.52. Prior to assessing mean gray values (MGV), the combined images were transferred to RGB stack images. MGV of both green and blue channels were calculated.

### Principal Component Analysis

For in-depth analysis of endometrium and endometriosis-derived spectral signatures, principal component analysis (PCA) was performed with the Unscrambler software (Unscrambler X10.5, CAMO, Oslo, Norway) on 200 extracted single spectra per Raman image from nuclei COL I masks. Briefly, PCA is a multivariate data analysis tool reducing the dimensionality of a set of spectral data on a vector-based approach, in which each vector, so-called principal component (PC), describes a variation in the spectra. Plotting PC values against each other visualizes a correlation or separation of two or more data sets.

### Deconvolution

For in-depth analysis of protein secondary structures, spectral deconvolution on the amide I region (1580-1720 cm^-1^) was performed on averaged COL I Raman spectra as described previously (24).

### Neural Network Classifier

Sample classification was undertaken by using a neural network from the open-source Keras and Tensorflow API (Google Brain). Information about the model architecture can be found in the supplementary methods. It was aimed to classify the data into one of the two classes: *control* and *diseased*. For classification, one dataset including both nuclei and COL I spectra was evaluated. The dataset consisted of 740 features (spectral range from 600-1800 cm^−1^ with a sampling interval of 4 cm^−1^ for both nuclei and COL I spectra) and 600 spectra per patient. The performance of CNN classification was improved by structure modification and hyperparameter tuning of the batch size, the number of epochs, the number of hidden units, the optimizer as well as the learning rate.

### Statistical Analysis

Statistical comparisons were performed from a minimum of three independent patients per menstrual cycle phase. Statistical analysis was performed using GraphPad Prism version 9.00 for Windows (GraphPad Software). Results are shown throughout the entire manuscript as mean ± standard deviation. Statistical significance was determined by one-way ANOVA or two-way ANOVA for multiple comparisons and grouped comparisons. Values of *p* < 0.05 were considered significant (*\*p* < 0.05; *\*\*p* < 0.01; *\*\*\*p* < 0.001). All *n*-numbers, applied tests and corresponding significance for each result are listed in the figure legends.

## Results

### Histological staining confirms the presence of characteristic glands in endometrial lesions and endometrium tissue

H&E and CD10 immunohistochemical staining were employed to identify endometrial gland positions in tissue sections. Samples from all phases of menstrual cycle were investigated. In H&E stains, extracellular matrix, cytoplasm (pink) and nuclei (blue) were visualized. Endometrium glands in control tissues in proliferative phase showed a typical single-layer high prismatic morphology (Figure 1b). Control tissue in the luteal phase showed a typical saw-blade shaped morphology with extended lumina. The CD10 staining confirmed the presence of endometrial stroma cells. Stroma around glands appeared brown, while nuclei of glands were stained in blue (Figure 1b). In dependence of the orientation of the tissue during sample processing into cryo-sections, glands appeared in sagittal shape. Using the H&E and CD10 staining as reference, gland positions were localized in consecutive, non-stained tissue section and investigated by RMS.

### Raman imaging enables marker-independent visualization of tissue structures

To characterize the biomolecular composition of glands and surrounding ECM, non-destructive and marker-independent Raman imaging and multivariate data analysis were implemented on control endometrium and endometriosis derived from LiS. Raman scans of glands were analyzed by true component analysis (TCA) and enabled to identify and localize major tissue structures (Figure 1c). Based on the identified Raman spectra, TCA allowed visualizing the glands as false color-coded intensity heatmaps in which each color represents the best fit to one of the identified molecular signatures. Raman spectra and morphology of structures visualized by TCA were applied to assign tissue structures to their biological origin which were nuclei (blue), lipids (yellow), collagen I (COL I, green) and endoplasmic reticulum (ER, pink). In endometriosis tissue sections, additionally to the spectral components found in control sections a further fiber-like structure was found (red), which was attributed to elastin as validated by resorcin-fuchsin staining (Supplementary Fig. S1) and literature reference spectra (22). Nuclei were identified by their typical peaks related to the phosphate stretching of the backbone of DNA located at 798 and 1096 cm^-1^ (27). Collagens were characterized by peaks at 855 and 936 cm^-1^ indicative for proline (28). The collagen component demonstrated a fibrillar distribution in the stroma. Lipids were identified by peaks located at 1442 and 1775 cm^-1^ explaining methylene groups and C=C vibrations in fatty acids (29, 30). Lipids are in close proximity to gland positions in both endometrium and endometriosis. Table 1 gives an overview of the identified Raman shifts and their corresponding biological explanation.

**Table 1.** Biological assignment of the most relevant wavenumbers.

| Wavenumber [cm<sup>-1</sup>] | Biological origin | Wavenumber [cm<sup>-1</sup>] | Biological origin |
| --- | --- | --- | --- |
| 676 | Guanine(36) | 1370 | CH<sub>3</sub>(34, 41) |
| 755 | Tryptophan(33) | 1397 | CH<sub>2</sub>(28) |
| 791 | PO<sub>2</sub><sup>-</sup> (27) | 1442 | CH<sub>2</sub> |
| 798 | PO<sub>2</sub><sup>-</sup> (27) | 1443 | CH<sub>3</sub>(58) |
| 855 | Proline(28) | 1494 | Guanine(59) |
| 936 | Proline(28) | 1486 | Adenine, Guanine(31) |
| 1091 | PO<sub>2</sub><sup>-</sup> (60) | 1552 | Amide II(61) |
| 1101 | PO<sub>2</sub><sup>-</sup>(31) | 1557 | Tryptophan (28) |
| 1163 | Tyrosine(28) | 1592 | C=C(61) |
| 1235 | Amide III (31) | 1628 | C=C(62) |
| 1291 | Cytosine(38) | 1666 | Amide I(61) |
| 1341 | Guanine(27) | 1775 | C=C(30) |
| 1362 | Cytosine(39) |  |  |

### Picrosirius-red staining identifies differences in collagen fiber density and orientation

To investigate collagen fibers more specifically, polarized light images of picrosirius red (PSR) stained tissue sections were obtained according to their birefringent characteristics. PSR images of endometrium and endometriosis were obtained for all menstrual cycle phases (Figure 2a). Regardless of the menstrual cycle phases, the quantification of the presence of dense or mature (red/orange) and thinner (yellow, green) fibers showed statistically significant differences between endometrium and endometriosis from LiS (Figure 2b). In endometrium, less mature and more immature collagen fibers were found in comparison to endometriosis tissues. This overall trend between endometrium and endometriosis was observed regardless of the menstrual cycle phases. However, for endometrium a decrease in red-polarized contribution is observed over the course of the menstruation cycle. While in PP more green fibers were present, OP and SP of endometrium showed an increase in red fibers (two-way ANOVA, *p* < 0.05). In contrast, no changes were observed across the menstrual cycle phases in endometriosis (two-way ANOVA, *p* > 0.05). Furthermore, quantification of collagen fiber alignment was performed by coherency analysis indicating the overall percentage of collagens that are parallelly aligned (Figure 2c). Non-statistically significant increases in coherency were found in endometriosis across all menstrual cycle phases compared to endometrium (two-way ANOVA, *p* > 0.05).

**Figure 2.**
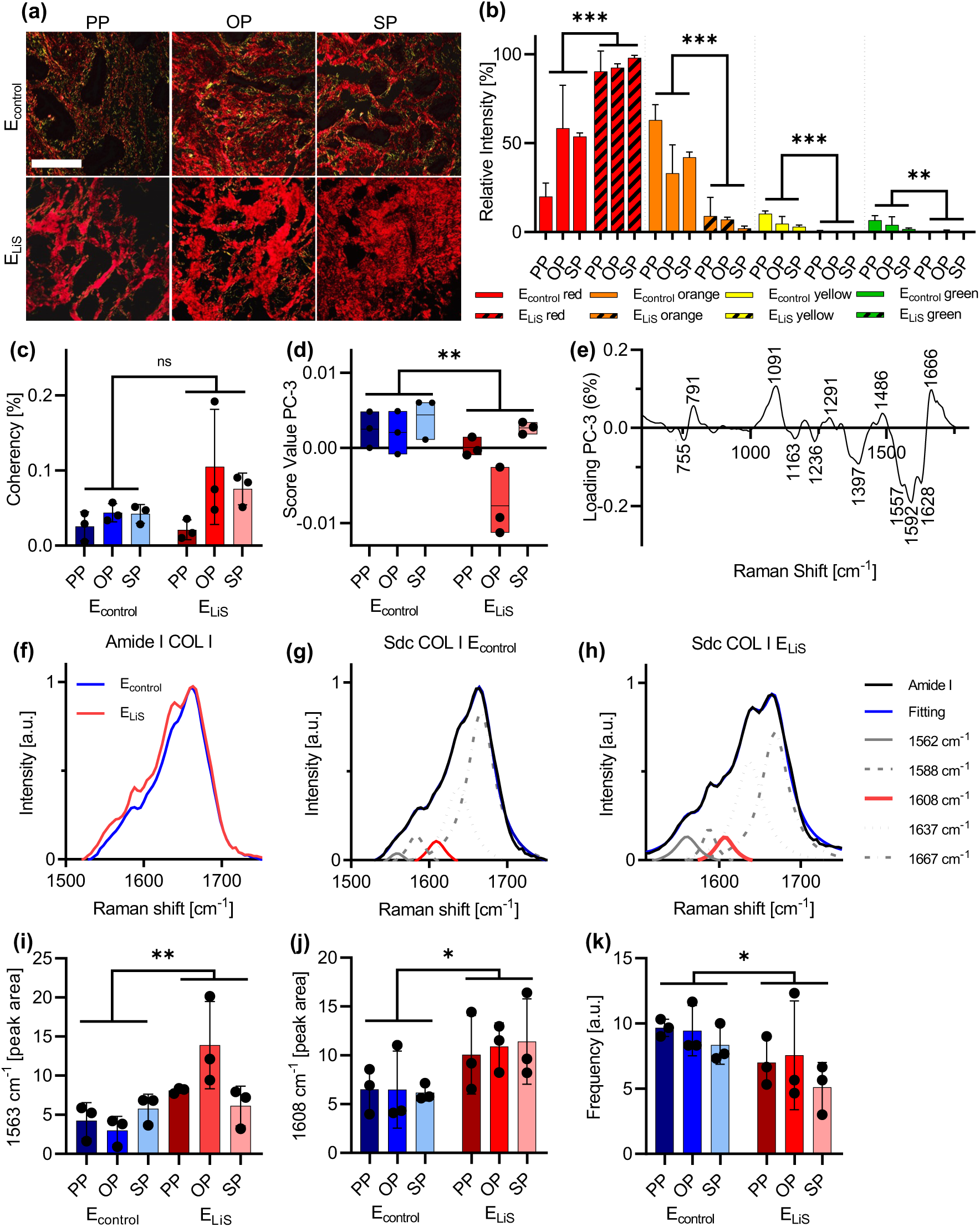
Collagen analysis by picrosirius red (PSR) staining and RMS reveal fibrotic changes in endometriosis. **(a)** PSR staining of endometrium *in loco typico* and endometriosis from LiS. Scale bar equals 100 µm. **(b)** Quantification of collagen fiber density using PSR color scales displays differences in collagen fiber structure in endometrium compared to endometriosis. **(c)** Fiber alignment of collagen fibers display a trend towards more parallel aligned fibers in endometriosis. **(d)** PCA score value analysis of COL I Raman spectra and **(e)** corresponding PC-3 loadings plot. **(f)** Amide I band of averaged COL I spectra from endometrium (blue) and endometriosis (red) were subjected to spectral deconvolution: **(g)** endometrium; **(h)** endometriosis. Peak area of subpeaks at **(i)** 1563 cm^-1^ and **(j)** 1608 cm^-1^ show differences between endometrium and endometriosis. **(k)** Frequency of modes from filter image ratio at 1608 cm^-1^ normalized by amide I maximum. Data are presented as a mean ± SD. Statistical differences between the groups were determined by two-way ANOVA (\**p* < 0.05, \*\**p* < 0.01, \*\*\**p* < 0.001, ns: not significant); *n*=3. E_control_: endometrium; E_LIS_: endometriosis of ligamentum sacrouterinum.

### COL I signatures differ between endometrium and endometriosis

In addition to image-based histopathological characterization of the tissues, in-depth analysis of the underlying Raman spectra allows for molecular-sensitive analysis of tissue structures. For submolecular analysis, principal component analysis (PCA) was performed on extracted COL I spectra (fingerprint range 400-180 cm^-1^). Despite impacts of the cycle phase in endometriosis derived collagens, significant differences were identified in COL I features between endometrium and endometriosis. Score value analysis of PC-3 showed decreased score values of endometriosis from LiS across all menstrual cycle phases (two-way ANOVA, *p* < 0.01) (Figure 2d). The loading plots identified prominent Raman peaks accounting for the separation (Figure 2e). Based on the loading of PC-3, increased spectral intensities correlating to endometrium were found at 1666 cm^-1^, representative for changes in amide I. The increased signals at 791 cm^-1^ and 1091 cm^-1^ in endometrium are attributed to DNA backbone vibrations (27), while the peaks at 1291 cm^-1^ and 1486 cm^-1^ are assigned to DNA bases (31, 32). The most prominent peaks in correlation to endometriosis were depicted at 1592 and 1628 cm^-1^, related to C=C bonds. Additional peaks assigned to the separation of endometriosis were located at 1397 cm^-1^, representative for CH_2_ deformations (33), 1236 cm^-1^ corresponding to changes in amide III (31) and 755 cm^-1^, 1163 cm^-1^ and 1557 cm^-1^ assigned to tyrosine and tryptophan (33, 34).

### Spectral Deconvolution Identifies Fibrotic COL I Alterations in Endometriosis

Continuing the analysis of COL I fibers from endometrium and endometriosis, Raman maps containing solely COL I spectra were extracted from large area scans and averaged. These Raman spectra were cropped to the amide I region (1520 - 1780 cm^-1^) that was utilized for the analysis of protein secondary structures. Spectral deconvolution was utilized to investigate underlying peaks of the amide I band (Figure 2f-h). The secondary structures of collagens are mainly α-like helices, β-sheets, β-turns, and random coils (disordered). The number and location of sub bands were selected by the shape of the amide I peak in COL I and as described previously (24). Comparison of the area under the curve at 1563 cm^-1^ (two-way ANOVA, *p* < 0.01(Figure 2i) as well as 1608 cm^-1^ (two-way ANOVA, *p* < 0.05) (Figure 2j) showed statistically significant changes between endometrium and endometriosis across all menstrual cycle phases. Peak areas at 1588 cm^-1^ (two-way ANOVA, *p* > 0.05) as well as 1636 cm^-1^ sub bands (two-way ANOVA, *p* > 0.05) showed no separation between endometrium and endometriosis (Supplementary Fig. S2). In one of our previous studies, calculating peak ratios (in an image-based approach) has been demonstrated to be a robust fibrosis biomarker (24), which could also be confirmed for the endometriosis tissues (two-way ANOVA, *p* < 0.05) (Figure 2k).

### PCA identifies differences in nuclei between endometrium and endometriosis

In addition to analysis of collagen structures, Raman spectra of nuclei from endometrial glands were extracted and analyzed by PCA, resulting in significant differences in PC-1 (two-way ANOVA, *p* < 0.001) (Figure 3a). Particularly pronounced peaks in the loadings (Figure 3b) separating nuclei features from endometrium and endometriosis were located at 794 and 1101 cm^-1^, indicating the phosphate backbone in DNA (35). Additionally, shifts at 676, 1341 and 1494 cm^-1^ corresponding to the DNA base guanine (36, 37), were observed in endometrium features. In endometriosis-derived spectra, increased intensities were found for amide III and amide II bands (1239 and 1552 cm^-1^) (31), cytosine (1285 cm^-1^ and 1362 cm^-1^) (38, 39) as well as methyl-groups (1443 cm^-1^) (23, 40).

**Figure 3.**
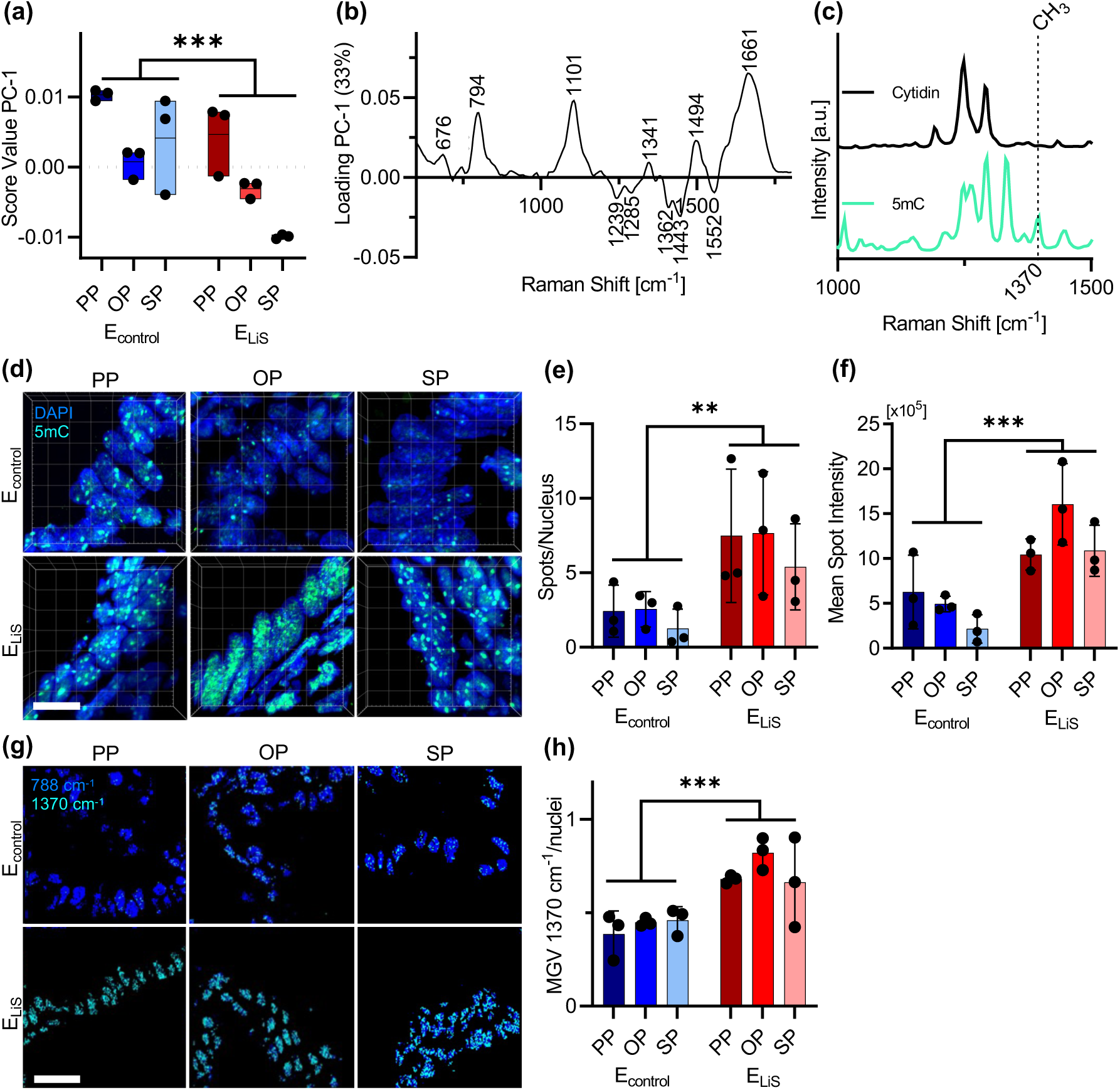
RMS and 5mC staining identify alterations in nuclei of endometriosis. **(a)** PCA of nuclei spectra indicate differences between endometrium and endometriosis. **(b)** Corresponding loadings plot. **(c)** Raman spectra of isolated cytidine and 5mC reference substances. **(d)** 5mC immunofluorescence staining (green) of endometrium and endometriosis. Nuclei: blue; Scale bar equals 10 µm. **(e)** Mean spot intensity of 5mC foci and **(f)** 5mC foci per nucleus were determined. **(g)** Overlay of Raman filter images at 1370 cm^-1^ (green) and 788 cm^-1^ (blue). Scale bar equals 30 µm. **(h)** An increase in 1370 cm^-1^ assigned mean gray value intensity (MGV) per nucleus is found in endometriosis. Data are presented as a mean ± SD. Statistical differences between the groups were determined by two-way ANOVA (\**p* < 0.05, \*\**p* < 0.01, \*\*\**p* < 0.001); *n*=3.

### 5mC staining demonstrates epigenetic influences in endometriosis

The differences observed in the Raman analysis of the nuclei, especially for peaks representative for methyl-groups and cytosine, encouraged to further elaborate on methylation patterns of the tissues. Subsequently, our objective was to determine epigenetic foci in Raman images of endometrium and endometriosis. In order to identify spectral bands indicative for epigenetic methylation, reference spectra of both cytidine and 5mC were recorded (Figure 3c). Comparison of both spectra revealed a peak assigned to CH_3_ at 1370 cm^-1^ only detected in 5mC (41, 42). IF staining of 5mC was utilized to identify epigenetic 5mC foci within nuclei (Figure 3d). Three-dimensional fluorescence images were analyzed to determine foci counts (two-way ANOVA, *p* < 0.01) and intensities (two-way ANOVA, *p* < 0.001), which were both found to be increased in endometriosis (Figure 3e,f). According to the generated 5mC reference spectra, sum-filter images at 1370 cm^-1^ were generated and combined with nuclei heatmaps recorded at 788 cm^-1^ (Figure 3g). Raman-based 5mC mean gray value (MGV) was increased in nuclei of glands in endometriosis (two-way ANOVA, *p* < 0.001) (Figure 3h).

### Fibrotic and epigenetic features discriminate endometriosis and surrounding tissue

RMS integrated in an endoscopic system offers the great perspective of being utilized intraoperatively to classify diseased tissue sites during a surgery without the need for biopsies and conventional histopathological staining. To elucidate the potential of RMS to identify endometriosis from surrounding tissues, glands from the LiS and a second ectopic position – SRV – were compared to healthy peritoneum. Therefore, the established analyses and extracted spectral features for COLI and 5mC were additionally performed on RMS data of peritoneum (P_control_) and endometriosis from SRV (E_SRV_) (Fig. 4a). The quantification of PSR color scales revealed similarities between endometriosis of LiS and SRV as well as peritoneum (one-way ANOVA, *p* > 0.05) (Fig. 4b). Overall, more mature fibers were detected in endometriosis and peritoneum compared to endometrium (one-way ANOVA, *p* < 0.001). Coherency analysis of collagen fibers (Fig. 4c) displayed no statistically significant differences between endometrium, peritoneum and endometriosis (one-way ANOVA, *p* > 0.05). However, a tendency towards more parallel aligned collagen fibers was visible by increased coherency in endometriosis. In contrast, spectral deconvolution of the amide I band of extracted COL I spectra revealed significant increases at peak areas of the 1608 cm^-1^ subpeak between control peritoneum and endometriosis of both LiS (one-way ANOVA, *p* < 0.01) and SRV (one-way ANOVA, *p* < 0.05) (Fig. 4d). Comparison within endometriosis showed no significant difference (one-way ANOVA, *p* > 0.05). Similar results were found in the analysis of the mode of filter images at 1608 cm^-1^ normalized by the amide I maximum in COL I spectra (Fig. 4e). Assessment of epigenetic alterations by 5mC foci demonstrated a significant discrimination between nuclei in glands of endometriosis from the LiS (one-way ANOVA, *p* < 0.01; *p* < 0.05; *p* < 0.01) and SRV (one-way ANOVA, *p* > 0.05; *p* < 0.05; *p* > 0.05) and peritoneum tissue for both IF as well as RMS-based localization (Fig. 4 f-h).

**Figure 4.**
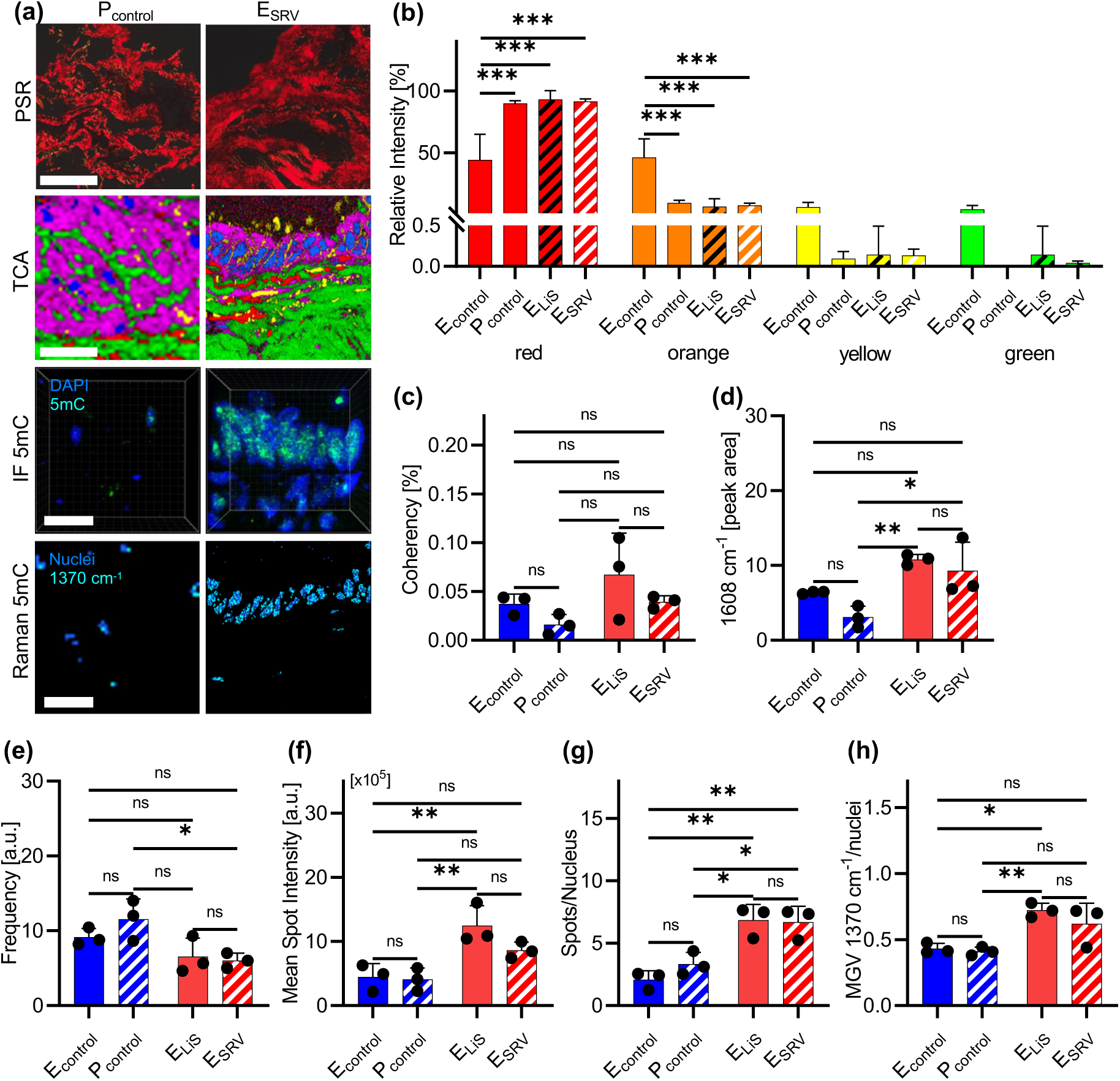
Fibrotic and epigenetic spectral biomarkers enable a discrimination between endometriosis and peritoneum. **(a)** Representative images of PSR, TCA, 5mC IF and combined Raman images at 1370 and 788 cm^-1^ of control peritoneum (P_control_) and endometriosis from the SRV (E_SRV_). Scale bars equal 100 µm (PSR), 20 µm (TCA), 10 µm (5mC IF) and 30 µm (5mC Raman). **(b)** PSR displays no statistically significant differences in collagen maturity/thickness between peritoneum and endometriosis from both LiS and SRV. (c) Fiber coherency of collagen fibers show no differences between peritoneum and endometriosis from both LiS and SRV. **(d**) Spectral deconvolution of the amide I region of averaged COL I spectra allow discrimination between peritoneum and endometriosis from both LiS and SRV. **(e)** Frequency of modes from filter image ratio at 1608 cm^-1^ normalized by amide I maximum of COL I. **(f)** Mean spot intensitiy from 5mC IF staining. **(g)** Spots per nucleus based on 5mC IF staining. **(h)** MGV analysis from Raman filter images at 1370 cm^-1^ normalized by whole nuclei signal. Quantitative data were presented as a mean ± SD. Statistical differences between the groups were determined by ordinary one-way ANOVA (\**p* < 0.05, \*\**p* < 0.01, \*\*\**p* < 0.001, ns; not significant); *n*=3.

### Neuronal Networks classify Raman signatures of endometriosis and peritoneum

To discuss the potential clinical suitability of Raman spectroscopy, the Raman spectra of nuclei and COL I of control peritoneum were compared to endometriosis from LiS and SRV and classified using a convolutional neural network (CNN). Applying a training-validation-test set split of 60-20-20 to the CNN lead to a test set error of 0.15% and test set loss of 0.64% at an accuracy of 98.9% with a validation loss of 0.64% and validation accuracy of 98.4% (Supplementary Table S2). The corresponding training set accuracy and test set accuracy as well as training set loss and test set loss across all epochs is displayed in Figure (Supplementary Fig. S3). Applying the trained CNN to the unknown test dataset yielded a prediction accuracy of 98.5%, sensitivity of 98.6%, and a specificity of 98.3%.

## Discussion

This study used non-invasive, label-free Raman spectroscopy to analyze the peritoneum, endometrium, and endometriosis of the LiS and the SRV. The results suggest that epigenetic and fibrotic factors may play a role in the development of endometriosis, as demonstrated through Raman measurements, IF imaging of epigenetic 5mC staining, PSR staining, and a novel Raman biomarker for fibrotic COL I analysis.

Collagen, the most prevalent and frequently modified constituent of the extracellular matrix (ECM), is fundamental for tissue formation and is linked to various diseases. Accurate control of ECM synthesis and remodeling is critical for human well-being. Aberrations in ECM remodeling can impact the progression of different pathological conditions, influencing disease course. Fibrosis is characterized by excessive ECM production with consecutive tissue stiffening, seen in cancer and skin ailments (43). Histological PSR staining was used for determining collagen fiber status. Despite controversial interpretations, it is generally reported that the color of stained collagen fibers can indicate a certain degree of maturity, which in turn correlates with birefringence properties (44). The PSR results showed that the endometrium’s functional layer synthesized new collagen fibers (yellow/green) during the proliferative phase. These matured (red/orange) during ovulation and the secretory phase, with loss during menstruation. In contrast, the stroma of endometriosis exhibited thicker, more mature collagen fibers. Due to their atypical location, the degree of maturity of the collagen fibers did not change during the menstrual cycle, and neither detachment nor new formation occurred. This suggests a distinct fibrotic ECM remodeling process in endometriosis. Fibrotic ECM remodeling is associated not only to collagen maturation but also to collagen fiber alignment. The parallel alignment of myofibroblasts, essential for collagen synthesis, is observed in fibrotic phenotypes *in vitro* and *in vivo*, resulting in parallel-aligned collagen fibers (24, 45) with concomitant increase in tissue stiffness (46).

The review by Vissers et al. also suggested that fibrosis should be included in the histopathological definition of endometriosis. Fibrosis in endometriosis is accompanied by processes such as epithelial-to-mesenchymal transition (EMT), fibroblast-to-myofibroblast transdifferentiation (FMT), and smooth-muscle-metaplasia (SMM), which lead to the formation of myofibroblasts. These myofibroblasts are the main producers of ECM in endometriosis and remain persistently active, continuously producing matrix proteins, which can in term lead to a loss of organ function (47) RMS further elucidated the molecular changes of COL I in the endometrium and endometriosis. The differences are mainly due to variations in C=C bonds and CH_2_ groups, accompanied by changes in amides I, and III as well as different proportions of tyrosine and tryptophan. These results indicate a change in the secondary structure of COL I. In a previous study we analyzed fibrotic diseases in various human organs and showed that fibrotic COL I alteration is marked by an increased peak area 1608 cm^-1^ in the amide I band of COL I Raman spectra. Application of this analytical tool showed evidence of the fibrotic COL I modification in endometriosis compared to endometrium as well as peritoneum (24).

We aimed to link the fibrosis-related pathogenesis in endometriosis with alterations on the nuclear level. RMS allowed differentiation of endometriosis and endometrium based on shifts in the phosphate scaffold of DNA and changes in prominent DNA bases, indicative of an altered chromatin structure. While increased spectral signatures for guanine are observed in the endometrium, cytosine signals dominate in endometriosis. In addition, signals relating to increased CH_3_ content, as well as elevated spectral intensities for amide III and amide II are found in endometriosis., possibly indicating changes in the histone packing of DNA. To investigate the increased levels of CH_3_ and cytosine in endometriosis we analyzed the tissues by their epigenetic 5mC profile. Both IF staining and Raman spectroscopy epigenetic analysis showed an increase in methylation and 5mC foci in endometriosis. These results suggest an alteration in chromatin packing affecting normal cellular activity. Epigenetic changes in methylation are reflected in lower proliferative activity of endometriosis compared with endometrium according to Ki67 staining (Supplementary Fig. S2).

In a review by Marquardt et al., several studies were analyzed that describe changes in histone deacetylases (HDAC) and acetyltransferases in endometriosis (48). In a separate study, Marquardt et al. observed a significantly reduced concentration of the HDAC3 protein in the eutopic endometrium of infertile women with endometriosis compared to control subjects (48). This change was associated with an increased expression of type I collagen in the endometrium - regardless of the cycle phase - and is linked to the development of fibrosis (48). They also hypothesized that an aggravation of inflammation in endometriosis lesions is caused by the upregulation of HDAC1 (48).

DNA methylation is one of the most thoroughly investigated and best characterized epigenetic processes (48). Numerous studies have reported altered DNA methylation patterns in endometriosis. In a recent study, epigenetic DNA methylation patterns were associated with the Heat Shock protein 47 (HSP47) gene expression (49). HSP47 has been reported to mediate TGFβ1-induced collagen deposition leading to fibrosis (45, 50). This finding agrees well with the increased peak area at the spectral position of 1608 cm^-1^ of COL I, which is associated with increased phenylalanine content found in COL I fibers increasing the binding affinity of HSP47 ultimately leading to fibrotic COL I alteration (24, 51, 52).

To further evaluate the robustness for assessing fibrotic and epigenetic alteration with Raman spectroscopy, we examined additional sites of endometriosis i.e., the SRV as well as the peritoneum. The latter served as a more realistic control, since endometriosis, other than adenomyosis uteri, is not localized in the endometrium. While histological PSR analysis was not sufficient to determine endometriosis from peritoneum, as the connective tissue is largely composed of mature collagen fibers, spectral deconvolution of COL I Raman spectra allowed to discriminate the tissues. Moreover, epigenetic differences in nuclei were observable by RMS.

Until now, the detection of endometriosis that had to be solved mainly by laparoscopic surgery is still challenging. Since 2000, Raman endoscope systems suitable for in vivo disease detection have been developed and expanded to preclinical and clinical investigations (53, 54).

In future applications, such endoscopes could be used in everyday clinical practice. This necessitates an automatized real-time classification of gathered Raman data to support the surgeon’s decision making. Using a neural network-based classification of Raman data from nuclei and COL I from peritoneum and endometriosis from the LiS and the SRV, we provide evidence for successful identification of the disease in an ex vivo model. It should be noted that the data-driven classification approach presented in this study is highly sample-specific, and its results may not apply to endometriosis and other control tissue in general due to the small number of patients involved. To further evaluate the robustness of this method for the detection of endometriosis, experiments would need to be performed with multiple donors and endometriosis lesions at other tissue sites such as from genitalia interna (i.e., myometrium), genitalia externa (i.e., vagina, perineum) or extra genitalis (i.e., ureter, retroperitoneum, lung, brain).

With RMS, it appears possible to assess the severity and progression of endometriosis in endometriomas (55). Akyuz et al. based their analysis on the observed increase in DNA concentration, kynurenine, and pyrrole content, as well as the decreased levels of tryptophan (55). Other studies have also demonstrated that the combination of PCA with types of discriminant analyses can reveal tumor development in brain samples (56), breast tissue samples (57) and liver samples (21) at the molecular level.

This study demonstrated the potential of RMS as a diagnostic approach to characterize and identify endometriosis. We showed the capability of RMS to discern endometriosis beyond influences of the menstrual cycle phase. Fibrotic COL I alterations were identified in endometriosis tissues. Moreover, RMS was able to detect 5mC-related epigenetic differences between endometriosis and endometrium as well as peritoneum. In combination with neural network-based classification, accurate tissue discrimination is enabled. The established spectral signatures allow to target structural and epigenetic alterations in a non-destructive manner and, in the future, could serve as novel biomarkers not only for investigations of endometrial pathogenesis but also as diagnostic markers. The presented approach could evolve to offer pathologists a complementary tool for non-destructive, marker-independent, and automated diagnosis of endometriosis with the potential to be implemented minimally-invasive by usage of Raman endoscopic systems.

## Conclusion

Raman imaging was successfully implemented as marker-free approach for histopathological assessment of endometrial and endometriotic tissue. Major tissue structures of gland tissue were localized and visualized based on specific spectral signatures. Multivariate analysis of individual spectral clusters identified submolecular changes in collagen I and DNA methylations between healthy endometrium and endometriosis that were consistent throughout cycle phases. In the future, these features could serve as biomarkers to complement histopathological analyses as well as for intraoperative tissue diagnostics if combined with an endoscopic setup.

## CREDIT Statement

L.B., T.B. and J.M. designed and conducted the study. M.W., K.R. and B.K. provided clinical samples. S.L. and D.B. assisted in tissue processing and staining. H.B. conducted pathological examination of the tissues. L.B. and T.B. performed statistical analyses. L.B. established neural network models. L.B. and T.B. were responsible for data processing and graphing. L.B., T.B., S.D. and J.M. interpreted the data. L.B., T.B. and J.M. wrote the manuscript. S.B, K.S-L., J.M. and M.W. supervised the project. All authors reviewed the manuscript.

## Acknowledgements

The work was conducted within the framework of the Graduate School 2543/1 “Intraoperative Multi-Sensory Tissue-Differentiation in Oncology” funded by the German Research Foundation (DFG - Deutsche Forschungsgemeinschaft) and the ENDO-RELIEF project (BMBF, 01EJ2403B). Further funding was received by the Deutsche Forschungsgemeinschaft (INST 2388/64-1, INST 2388/33-1 and Germany’s Excellence Strategy, EXC 2180-390900677), the Ministry of Science, Research, and the Arts of Baden-Wuerttemberg (33-729.55-3/214 and SI-BW 01222-91) and the State Ministry of Baden-Wuerttemberg for Economic Affairs, Labour and Housing Construction (3-4332.62-NMI/65).

## Supplementary material

Supplementary data to this article can be found online.

