## Supplementary information for "Marker-independent imaging reveals a correlation of fibrotic and epigenetic alterations in endometriosis"

**Supplementary Tables**

**Table S1.** Patient ages per donor and menstrual cycle phase. PP: proliferative phase; OP: ovulation phase; SP: secretory phase.

| **Sample** | **Donor 1** | **Donor 2** | **Donor 3** |
| --- | --- | --- | --- |
| Endometrium PP | 41 | 40 | 49 |
| Endometrium OP | 38 | 45 | 42 |
| Endometrium SP | 37 | 44 | 42 |
| Endometriosis LiS PP | 31 | 41 | 36 |
| Endometriosis LiS OP | 40 | 29 | 38 |
| Endometriosis LiS SP | 31 | 43 | 40 |
| Endometriosis SRV | 31, PP | 41, PP | 32, OP |
| Peritoneum | 25 | 31 | 31 |

**Table S2.** Performance of CNN classification of nuclei and COL I Raman spectra from endometriosis from both LiS and SRV and peritoneum.

| **Training** | CNN |
| --- | --- |
| Test set error | 0.015 |
| Test set loss | 0.065 |
| Training accuracy | 0.984 |
| Validation loss | 0.064 |
| Validation accuracy | 0.985 |
| **Test** |  |
| Accuracy | 0.985 |
| Sensitivity | 0.985 |
| Specificity | 0.983 |

**Supplementary Methods**

**Detailed protocols of histological stains.**

### Hematoxylin-Eosin (H&E) Staining

Cryo-sections were washed twice with DPBS for 10 min and fixed with 4% para-formaldehyde (PFA) for 20 min. Mayer's hemalum solution (Roth, Karlsruhe, Germany) (8 min) was used as a stain for the nuclei and eosin (Roth) (3.5 min) served as the counterstain. Differentiation was reached by rinsing the section with distilled water. Sections were then dehydrated in an ascending ethanol series ending with 100% isopropanol and consecutive mounting with Isomount 2000 (VWR Chemicals, Radnor, USA).

### CD10 Staining

Cryosections were washed and fixed. Endogenous peroxidase activity was quenched with 1.5 ml 30% H_2_O_2_ (Sigma Aldrich, St. Louis, USA) in 50 ml methanol (Thermo Fisher Scientific, Waltham, USA) in the dark (20 minutes). Unspecific binding sites were blocked with horse serum (vector R.T.U Normal horse serum 2.5% 50 ml (Thermo Fisher Scientific) (30 min). After washing with DPBS^-^, anti-CD10 mouse IgG1 (1: 100, Leica Biosystems, Nußloch, Germany) was applied overnight at 4 °C. Sections were rinsed three times with DPBS^-^, then ImmPRESS™ Reagent anti-mouse IgG ready-to-use solution (Thermo Fisher Scientific) was applied (30 min). Sections were then incubated with ImmPACT™ DAB kit (Abcam, Cambridge, GB) under continuous light microscope control. After washing with DPBS^-^, nuclei were stained with hematoxylin QS (Vector Laboratories, Newark, USA). Finally, sections were mounted with Dako Faramount aqueous mounting medium (Agilent, Santa Clara, USA).

### Ki67 Staining

Cryo-sections were washed and fixed. Slides were permeabilized with 1% Triton X-100 (Sigma Aldrich) for 30 min. After 10 min of washing with PBS, unspecific binding sites were blocked with ImmPRESS horse serum (Thermo Fisher Scientific) for 30 min. Ki67 primary antibody (1:1000, Abcam) was applied and incubated overnight at 4 °C. Cryo-sections were washed three times with 0.01% Tween in PBS before applying ImmPRESS-AP Horse Anti-Rabbit IgG Polymer Kit+Alkaline Phosphatase Polymer Reagent (Thermo Fisher Scientific) for 30 min. After washing three times with 0.01% Tween in PBS ImmPACT Vector Red Substrate Kit (Thermo Fisher Scientific) was applied for 15 min in the dark. Finally, sections were mounted with Dako Faramount aqueous mounting medium.

*Resorcin Fuchsin Staining*

Cryo-sections were prepared as usual. Sections were placed in resorcinol-fuchsin staining solution (Sigma Aldrich) for 15 minutes. Nuclei were stained with red aluminum sulfate for 8 minutes. Sections were then rinsed with water for 1 minute. Differentiation was performed with 0.5% HCl in ethanol. Finally, the slides were mounted with Dako Faramount aqueous mounting medium.

### Picrosirius Red Staining

Collagen maturity and directionality analyses were performed via picrosirius red staining. Firstly, Weigert's hematoxylin was used to stain nuclei of cryosections for 8 minutes and washed with tap water for 10 minutes. The sections were then treated with 0.1% picrosirius red solution (Morphisto, Frankfurt/Main, Germany) for 60 minutes. After the treatment, the tissues were washed with 0.5% acetic acid and 100% ethanol. The picrosirius red stained sections were imaged by polarized light microscopy (Axio Observer, Zeiss Microscopy GmbH, Oberkochen, Germany) at 40x magnification followed by ImageJ (Fiji version 2.0.0) processing. The images were transferred to RGB colors. In order to acquire the area percentages of red, orange (mature collagens), yellow and green (immature collagens) signals, the thresholds were adjusted as follows: Red (1-13, 230-256), orange (14-25), yellow (26-52) and green (53-110). For morphological assessment of collagen fibers, we utilized methods based on curvelet transform and Fourier transformation. CURVE Align, an open-source software package, was used to calculate alignment of collagen fibers. Alignment was represented on a scale from 0–1, where 1 displayed the highest degree of parallelism.

### 3D-image analysis

5mC and DAPI channels were analyzed separately with Imaris 9.7.2. The DAPI channel was processed individually for optimal visibility and contrast as it serves for general identification of nuclei. First, image layers negative for DAPI were truncated. Layers were then intensity normalized to 1. For calculation of nuclei volumes, the “Surfaces” tool was utilized to create artificial solid objects of a specified gray value range. Gaussian smoothing was set to 10^-4^ µm. To separate individual nuclei, “Region growing” was set to 0.1 µm. A threshold of a minimum of 10 voxels was applied for final filtering. Non-glandular nuclei volumes were deleted manually. The remaining surfaces were unified. To determine the number of glandular nuclei in a z-stack, “ClearView-GPU Deconvolution” with standard parameters utilizing the “Robust” algorithm with a maximum of 25 iterations was utilized. Resolved nuclei were independently counted manually three times. For 5mC analysis, histogram values were set between 1500 and 15000. First, control stains with no primary antibody were analyzed to determine the background noise. The “Threshold cutoff” was set to an intensity of 1500. For analysis of intensity of 5mC within nuclei, previously calculated surfaces of glandular nuclei were utilized. In the statistics tab of the created surface mean intensity, summed intensity and volume within the surface of 5mC were acquired. To identify 5mC spots, the “Spots” function was utilized. “Classify Spots” as well as “Region Growing” was disabled for classification. The estimated spot size was set to 1 µm. “Background Object Subtraction” was enabled for spot detection. Created spots were filtered with a quality above 1500. All spots that were not distributed within the glandular nuclei surface were deleted.

### Neural network architecture

The neural network model had a fully-connected layer structure with 512-256-128-2 nodes, where the layer with 512 nodes was the input layer and the one with 2 nodes the output layer. The activation functions used were mainly ReLU (Rectified Linear Unit) and one softmax function in the last layer for probabilistic output values. Hyperparameter tuning was performed to identify optimal parameters for classification. The used hyperparameters were batch size: 128, epochs: 100, number of hidden units: 512, optimizer: Adam, learning rate: η=0.001. A training-validation-test split of 0.6-0.2-0.2 was applied. In our data assessment we compared dense neural networks with different amounts of layers as well as different amounts of dropouts in between the layers. The best classification results were obtained with a dropout of 75% after the first and after the second hidden unit.

**Supplementary Figures**

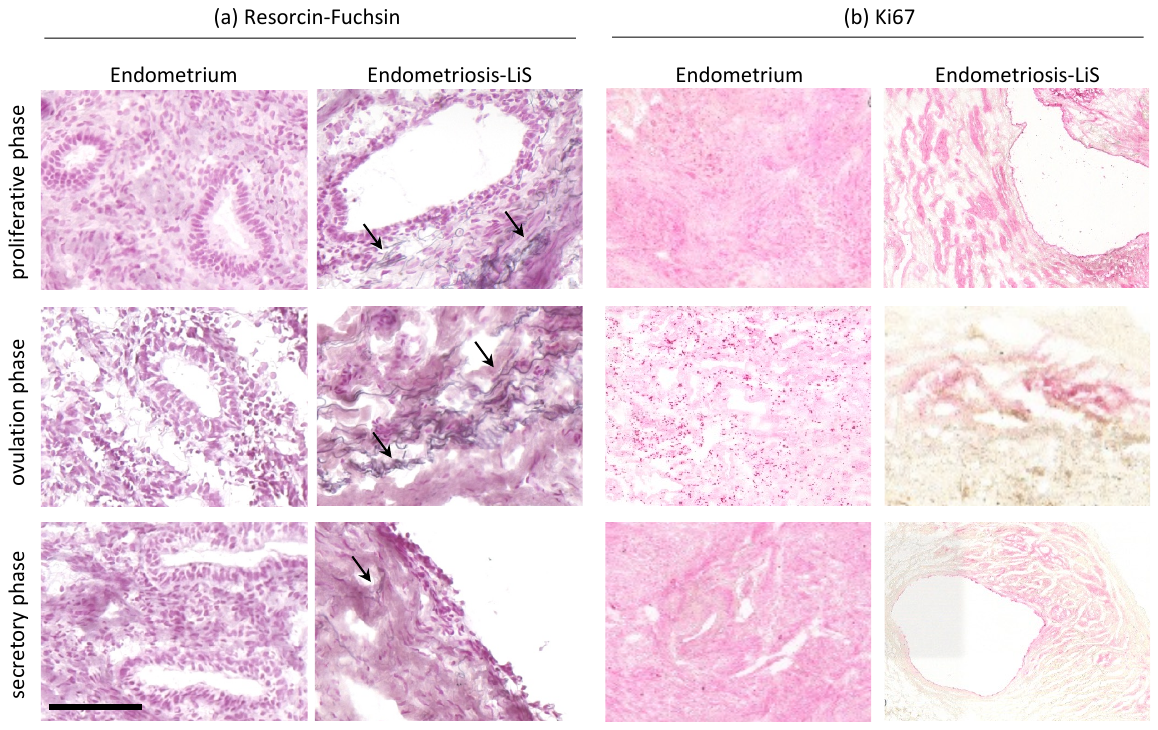

**Figure S1: Resorcin Fuchsin staining for detection of elastic fibers and Ki67 staining for identification of proliferation.** (a) Resorcin fuchsin staining of the endometrium and endometriosis of LiS in different menstrual cycle phases indicate presence of elastic fibers in endometriotic tissue (indicated by arrows). Scale bar equals 100 µm. (b) Ki67 staining of endometrium and endometriosis of LiS in different menstrual cycle phases. Proliferative cells are visible by pink staining. While endometrium shows staining in the whole tissue, in endometriosis, proliferation takes place near glands.

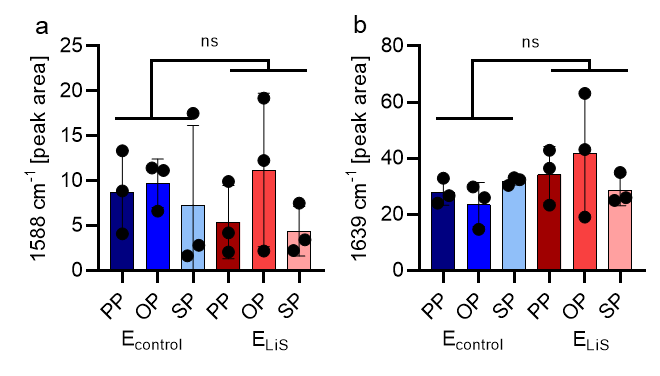

**Figure S2: Spectral deconvolution (Sdc) of the amide I region from averaged COL I Raman spectra display differences between endometrium (E_control_) and endometriosis from the LiS (E_LiS_).** Peak areas calculated based on spectral deconvolution of amide I area of averaged COL I spectra at (a) 1588 cm^-1^ and (b) 1639 cm^-1^ display no difference between endometrium and endometriosis from the LiS.

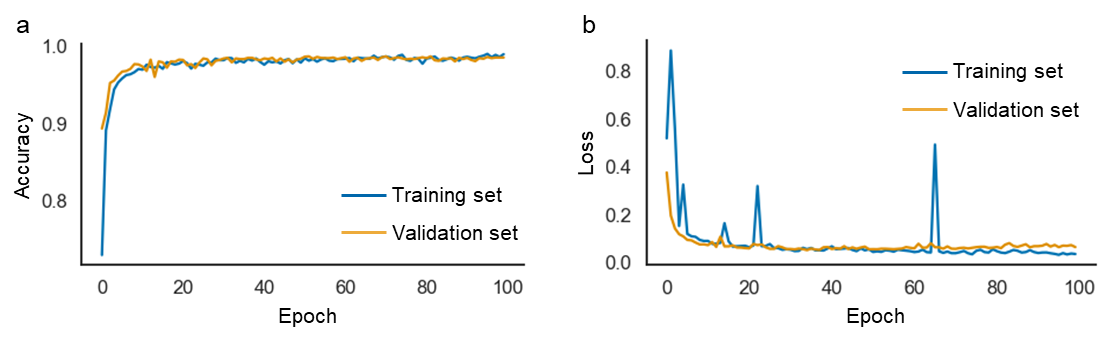

**Figure S3: Training and validation curves of CNN classification.** (a) Accuracy of training and validation set classifying endometriosis form both LiS and SRV and peritoneum based on combined COL I and nuclei Raman data. (b) Loss of training and validation set.
